# Closed-loop optimization of transcranial magnetic stimulation with electroencephalography feedback

**DOI:** 10.1101/2021.08.31.458148

**Authors:** Aino E. Tervo, Jaakko O. Nieminen, Pantelis Lioumis, Johanna Metsomaa, Victor H. Souza, Heikki Sinisalo, Matti Stenroos, Jukka Sarvas, Risto J. Ilmoniemi

## Abstract

**Background:** Transcranial magnetic stimulation (TMS) is widely used in brain research and treatment of various brain dysfunctions. However, the optimal way to target stimulation and administer TMS therapies, for example, where and in which electric field direction the stimuli should be given, is yet to be determined.

**Objective:** To develop an automated closed-loop system for adjusting TMS parameters (in this work, the stimulus orientation) online based on TMS-evoked brain activity measured with electroencephalography (EEG).

**Methods:** We developed an automated closed-loop TMS–EEG set-up. In this set-up, the stimulus parameters are electronically adjusted with multi-locus TMS. As a proof of concept, we developed an algorithm that automatically optimizes the stimulation orientation based on single-trial EEG responses. We applied the algorithm to determine the electric field orientation that maximizes the amplitude of the TMS– EEG responses. The validation of the algorithm was performed with six healthy volunteers, repeating the search twenty times for each subject.

**Results:** The validation demonstrated that the closed-loop control worked as desired despite the large variation in the single-trial EEG responses. We were often able to get close to the orientation that maximizes the EEG amplitude with only a few tens of pulses.

**Conclusion:** Optimizing stimulation with EEG feedback in a closed-loop manner is feasible and enables effective coupling to brain activity.

## Introduction

Transcranial magnetic stimulation (TMS) [1] has shown therapeutic promise in various brain disorders [2] with almost no side effects [3]. Rapidly repeated magnetic pulses can modulate neuronal activity and yield, for instance, anti-depressant effects and improved recovery from stroke [2]. Although TMS has been investigated for decades and applied in therapeutic use for several years, the optimal way to deliver TMS therapy in different clinical conditions and individuals is still unclear.

TMS-induced brain activation is produced by inducing a focused electric field (E-field) in the brain with a coil placed on the head. Targeting the stimulating E-field, i.e., selecting the location and orientation of its estimated focus on the cortex, is a key step in TMS. Other stimulus parameters include, for example, the stimulation intensity and the time between consecutive pulses, making the space of available TMS parameters huge. In TMS treatments and TMS studies investigating brain mechanisms, these parameters have been chosen in different ways. A common approach has been to observe and map functional responses to TMS, including motor-evoked potentials measured with electromyography when stimulating the motor cortex [1], subject-reported phosphenes or visual suppression upon stimulating the occipital cortex [4], and speech disruptions when perturbing language areas [5]. Stimulation locations have also been selected based on bony landmarks [6] or standard electroencephalography (EEG) electrode positions [7,8]. TMS targeting and mapping of functional responses got more precise with the introduction of neuronavigation systems [9,10], which guide and record the selection of the locations and orientations of the E-field focus based on individual structural brain scans.

Accurate TMS target selection is crucial: TMS– electromyography studies have shown that even subtle changes in the stimulation parameters may lead to large changes in the motor responses [11–14]. There are changes also in TMS-evoked potentials (TEPs) measured with EEG when the stimulation parameters change significantly [15–19], although detailed mappings of TEPs are lacking. Detailed TEP maps as a function of different stimulus parameters would increase understanding of the brain dynamics underlying TMS–EEG, which provides a direct measure of evoked brain activity. TMS–EEG is effective in investigating brain mechanisms and neuronal connections [20,21]; it has shown promise as a biomarker in predicting the outcome and monitoring the clinical efficacy of TMS or other neuromodulatory treatments [21–23].

Given the varying practices in TMS targeting and the sensitivity of the TMS responses to the stimulus parameters, it is unsurprising that the observed TMS effects are often different and difficult to interpret; sometimes they even appear contradictory [2,24,25]. To make TMS more effective and more comparable, the stimulation parameters should be adjusted with neurophysiological feedback, regardless of the cortical target region. This concerns not only the initial stimulation parameters selected at the beginning of a TMS session but also the real-time adjustment of the TMS parameters during treatment sessions to maximize the desired plastic effects. It has been suggested that TMS parameters could be adjusted with feedback from EEG [21,22,26], which provides a readout signal from superficial brain areas. Procedures for selecting initial stimulation parameters in TMS–EEG measurements by visually inspecting averaged EEG responses and manually adjusting the TMS settings have been applied, e.g., in Refs. [27–29] to acquire high-quality, artifact-free TEPs. There are, however, also other situations, such as selecting stimulus parameters for TMS treatments, that could benefit from EEG-based targeting. To make EEG-guided TMS practical and generally applicable, the analysis of EEG data and the adjustment of TMS parameters must be easy and fast—leaving automated closed-loop control [30] as the only feasible approach. Previously, the timing of TMS pulses has been automatically adjusted based on the phase of the pre-stimulus EEG [31,32]. Such methods are, however, considered only as state-dependent stimulation, not closed-loop stimulation that would utilize TMS responses in the process [33]. We hypothesized that TMS parameters could be automatically and adaptively adjusted based on the cortical effects of TMS as measured by EEG, even though TEPs exhibit high variation.

In this paper, we show that guiding TMS with online EEG responses is possible and present an automated algorithm for determining an optimal stimulation orientation based on TEPs. In our closed-loop set-up, the stimulation parameters are adjusted electronically with multi-locus TMS [14], which avoids conventionally required manual coil positioning by simultaneously operated overlapping coils (see Fig. *1*a). We show how TEPs depend on the stimulation orientation on the left pre-supplementary motor area (pre-SMA). Moreover, we demonstrate the functioning of the automated TMS–EEG targeting (Fig. *1*b) in the search of an optimal stimulation orientation to maximize the peak-to-peak amplitude of the early TEP deflections. The presented algorithm enables user-independent and individually optimized TMS targeting, especially outside the primary motor cortex. Furthermore, the possibility for automatic tuning of stimulus parameters with EEG feedback during TMS sessions has great potential for increasing the efficacy of TMS treatments.

**Figure 1.**
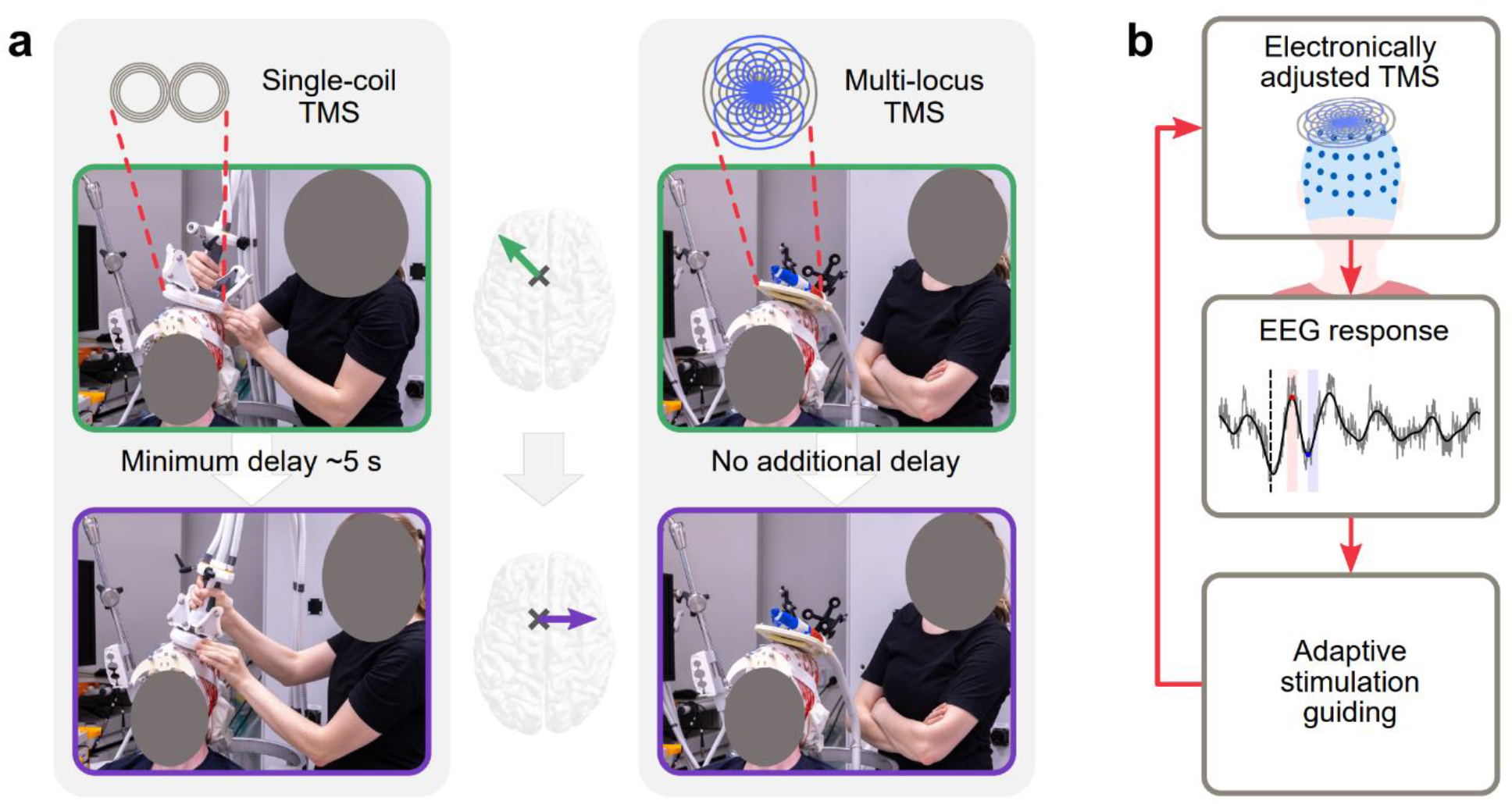
Benefit of multi-locus TMS in closed-loop TMS–EEG. **a** Instead of manual coil operation (left), multi-locus TMS (right) allows electronic adjustment of stimulus parameters (in this example, the stimulation orientation (middle)) with no delay. The transducer consists of two tailored overlapping coils (top right). **b** In automated TMS–EEG targeting, the evoked EEG responses are analysed in real-time and used to decide on the stimulation parameters for the next pulse in such a way that the optimal stimulation parameters are found with the least number of iterations. The stimulation parameters are effortlessly adjusted with multi-locus TMS. The loop is repeated until the optimal stimulation parameters are found.

## Methods

We performed two experiments. In Experiment 1, we systematically recorded TEPs as a function of stimulation orientation. Experiment 2 comprised repeated testing of the automated optimization of the stimulus orientation with EEG feedback. At the beginning of the TMS–EEG measurements, we performed a preparatory experiment comprising testing of the auditory noise masking and selection of the stimulation location and intensity.

Six healthy participants (2 males, 1 left-handed [34], aged 22–42) volunteered for the study. The participants were divided into two groups of three subjects. Group A had the preparatory experiment and Experiment 1 in a single session, and the corresponding data were used to tune the algorithm validated in Experiment 2 about 1–2 months later. For the Group B subjects, Experiment 2 was performed first to provide an independent validation of the algorithm, and Experiment 1 was carried out about one week later. Prior to the TMS–EEG experiments, the subjects underwent structural magnetic resonance imaging (MRI) with T1-, fat-suppressed T1-, and T2-weighted sequences (cubic voxel size 1 mm^3^). The study was accepted by the ethical committee of the Hospital District of Helsinki and Uusimaa and carried out in accordance with the Declaration of Helsinki. The subjects signed a consent form before the experiments.

### Measurement set-up for TMS–EEG

With our multi-locus TMS system [14] and a 2-coil transducer [35], we applied a monophasic magnetic pulse (60-μs rise time, 30-μs hold period, and 44-μs fall time [36]), producing a biphasic E-field in the cortex. The multi-channel TMS system allowed electronic adjustment of the stimulus orientation, here defined as the direction of the peak E-field computed on the cortex at 15-mm depth in a spherical head model of 85-mm radius. We attached a thin foam pad under the transducer to reduce the vibration of the electrodes and bone-conducted transmission of the stimulus sound. The transducer placement relative to the subject’s head was tracked with an eXimia NBS 3 neuronavigation system (Nexstim Plc, Finland), which during repeated stimulation helped to keep the transducer within 2 mm and 2° from the intended coil placement.

EEG signals were recorded with BrainAmp DC amplifiers (Brain Products GmbH, Germany). The signals were low-pass filtered with a 1000-Hz cut-off frequency and sampled at 5000 Hz. During TMS–EEG data acquisition, subjects were instructed to be relaxed, avoid swallowing, keep their sight fixated to a point, and when needed, blink preferably about 1 s after the pulses. To minimize the auditory responses evoked by the click sound of the TMS pulse, subjects were presented via earbuds auditory masking that contained white noise and randomly jittering click noise from a recorded coil click [37] (https://github.com/iTCf/TAAC). Subjects wore earmuffs to additionally attenuate the TMS click.

### Preparatory step: selecting fixed experimental parameters

Suitable auditory noise masking was determined by delivering TMS with the maximum intensity with the transducer a few centimetres above the head while the noise volume was increased until the subject did not hear the coil click. The effectiveness of the noise masking was tested by recording sets of 20 TEPs, which were evaluated with the help of a TMS–EEG data visualization tool presented and shared in Ref. [29] (https://github.com/iTCf/rt-TEP). This tool allowed cutting out the high-amplitude TMS pulse artifact within the first few milliseconds after the pulse and showing average-referenced signals to ease the visual inspection of the data. If auditory components were present, the noise masking volume was increased until the auditory components were negligible or the subject’s comfort limit or a 90-dB safety limit (measured with an SM20-A sound meter, Amprobe, USA, in a plastic ear canal) was reached.

The stimulation location (site of the estimated E-field maximum) on the left pre-SMA was placed over the superior frontal gyrus approximately 1–1.5 cm anterior to the vertical anterior commissure line [38]. The stimulation intensity was adjusted until in 20 trials, the average peak-to-peak amplitude of the deflections 15–50 ms after the pulse was 5–10 μV (similar approach as in Refs. [28,29]). We assessed the signal quality with the data visualization tool [29] with the induced peak E-field in the posterior– anterior and medial–lateral directions. If large stimulation artifacts were present in the electrodes close to the stimulation site, the location was changed a few millimetres to reduce the artifacts.

### Experiment 1: systematic mapping of orientation dependency of TEPs

In Experiment 1, 48 TMS pulses were delivered in each of the 36 orientations (with 10° steps) on the left pre-SMA, with the stimulus placement and intensity being as determined in the preparatory experiment. The 1728 pulses were divided into 12 blocks (a few minutes break between), each of them including four pulses in all 36 orientations in a pseudorandom order. At the beginning of each block, we gave one additional pulse in a random orientation; the corresponding data were excluded from the analysis. The interstimulus interval (ISI) was 2.4–2.7 s.

The EEG data were processed and analysed with Matlab scripts (version R2020b or newer; The MathWorks, Inc., USA). In each trial (−600...600 ms around the TMS pulses), the stimulation artifact in the time interval of −2...8 ms was removed and replaced by signals obtained by piecewise cubic interpolation. The signals were high-pass filtered with a third-order Butterworth filter (cut-off frequency 1 Hz) in the forward and backward directions. Bad trials containing eye blinks or an excessive amount of muscle activity were manually removed. The signals were baseline-corrected by subtracting the mean of the signal at −200...−10 ms from the whole trial. Then, we applied the source-estimate-utilizing noise-discarding (SOUND) algorithm [39,40] to diminish artifactual signal components, such as muscle activity. SOUND was applied separately for each pool of trials with the same stimulation orientation and with the tuning factor for the regularization parameter set to 0.1, channel C6 serving as a high-quality reference electrode, and the number of iterations being 10. In the minimum-norm estimation included in SOUND, the lead fields were based on the individual realistic head geometry. Structures of the head were segmented from fat-suppressed T1 and T2 MRIs with the SimNIBS *headreco* pipeline [41], followed by mesh downsampling and smoothing. The conductivity model consisted of scalp, skull, and intracranial volume, with conductivities of 0.33, 0.0066, and 0.33 S/m, respectively. The lead fields were computed with the boundary element method using a linear-collocation isolated-source approach [42,43] (https://github.com/MattiStenroos/hbf_lc_p) for source space on the grey–white matter boundary discretized to 25,000 dipoles normal to the surface of the cortex. The SOUND step was followed by low-pass filtering (cut-off frequency 45 Hz) and downsampling of the signals to 1000 Hz. Finally, the data were average referenced by subtracting the mean of all channels from each channel.

The peak-to-peak amplitude of the P20–N40 complex (subtraction of the signal minimum within the 35...45 ms interval from the signal maximum within the 15...25 ms interval) in channel FC1 was extracted separately for each trial (see an example TEP trial in Fig. *1*b with the corresponding peaks and time intervals highlighted). The dependence of the average P20–N40 amplitude as a function of the stimulus orientation was computed for the 36 stimulation orientations as an average of the responses within 30° (inclusive) from the computation point. Between the sampled orientations, the mean curve was generated by cubic interpolation. The standard deviation of the P20–N40 amplitudes was determined by first computing the variance over the single-trial P20–N40 amplitudes separately for each stimulus orientation, followed by averaging the variances and taking the square root. The SNR of the P20–N40 amplitude was determined as the ratio of the amplitude range of the mean curve over different stimulation orientations and the standard deviation of the single-trial P20–N40 amplitudes.

### Experiment 2: automated orientation search

In Experiment 2, we performed 20 times the automated search of the optimal stimulation orientation on the left pre-SMA with EEG feedback. In this experiment, we had 360 possible stimulation orientations separated with 1° steps; the ISI was in the range of 2–6 s. The stimulus placement and intensity were as determined in the preparatory experiment. The automated orientation search algorithm was based on our previously developed algorithm for optimizing TMS parameters with the motor response as the feedback signal to be maximized [44]. Here, we made that algorithm work for maximizing TEP amplitudes (see Fig. *1*b). Our adaptive search algorithm is founded on Bayesian optimization [45], which is an efficient approach to find a global optimum (e.g., maximum or minimum) of an unknown function. The EEG feature to be optimized was selected as the peak-to-peak amplitude of the P20–N40 complex in the FC1 electrode, as it showed orientation dependency consistently across the Group A data (Subjects 1–3) in Experiment 1.

The search process started by delivering a TMS pulse at a random E-field orientation followed by a pulse in the opposite orientation. Subsequently, we gave stimuli at orientations indicated by a guiding function called knowledge gradient, which was computed on a logarithmic scale for numerical accuracy [46]. The knowledge gradient suggests sampling points balancing high expected amplitude and large uncertainty so that the optimum is found with a minimal number of pulses. We modelled the dependence of the P20–N40 amplitude as a function of the stimulation orientation with Gaussian process regression, which estimates the underlying function by smoothly linking the neighbouring data points with the help of a covariance function [47]. We applied the following periodic (periodicity of 360°) covariance kernel function [47]: 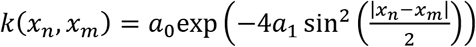, where *x*_*n*_ and *x*_*m*_ are the orientations for which the covariance is computed for, *a*_0_ determines the amplitude variance, and *a*_1_ the smoothness of the function to be fitted. The covariance parameters *a*_0_ and *a*_1_ along with the other parameters for the prior and the likelihood models were defined as by Tervo *et al*. [44]. The posterior mean curve (see examples in Fig. 3a,d), whose maximum indicated the estimated optimal stimulation orientation, was computed with a grid spacing of 0.25°. The minimum and the maximum number of samples in the search were set to 30 and 60, respectively. We considered the search converged when the estimated optimal stimulation orientation had not changed more than 5° during 10 consecutive iterations.

The EEG signals were transferred in real-time to another computer running our Matlab-based algorithm with the help of the code examples provided by Brain Products (https://www.brainproducts.com/downloads.php?kid=2&tab=5). The real-time processing of the EEG signals was similar to that in Experiment 1. However, we did not apply the SOUND algorithm, and the filtering was combined into a sixth-order band-pass Butterworth filter (1–45 Hz). The baseline correction (mean-computation interval −500...−10 ms) was applied as the second-to-last step before average referencing. If the signal range of a processed TEP in any channel exceeded 75 μV within the time interval of −500...500 ms, the trial was rejected and a new trial acquired (on average 1.8 rejected trials per search; subject-wise averages ranging from 0.1 to 5.6). However, for Subject 3, the rejection threshold was increased to 150 μV, as the data were excessively noisy (9.7 rejections per search on average).

The performance of the EEG-based orientation search algorithm was evaluated by comparing the search outcomes with the mean curve computed from the data measured in Experiment 1 (see Fig. 4a–f); the error of each search run was defined as the absolute difference of the optimized stimulation orientation and the closest maximum of the individual mean curve.

## Results

### Experiment 1: orientation dependency of the TMS–EEG responses

Figure 2a,d,e shows how the averaged TEPs of Subject 1 varied as a function of the E-field orientation (Fig. 2c). As expected, the largest early responses were evoked in the channels close to the stimulation site, and their amplitude varied with the stimulation direction.

**Figure 2.**
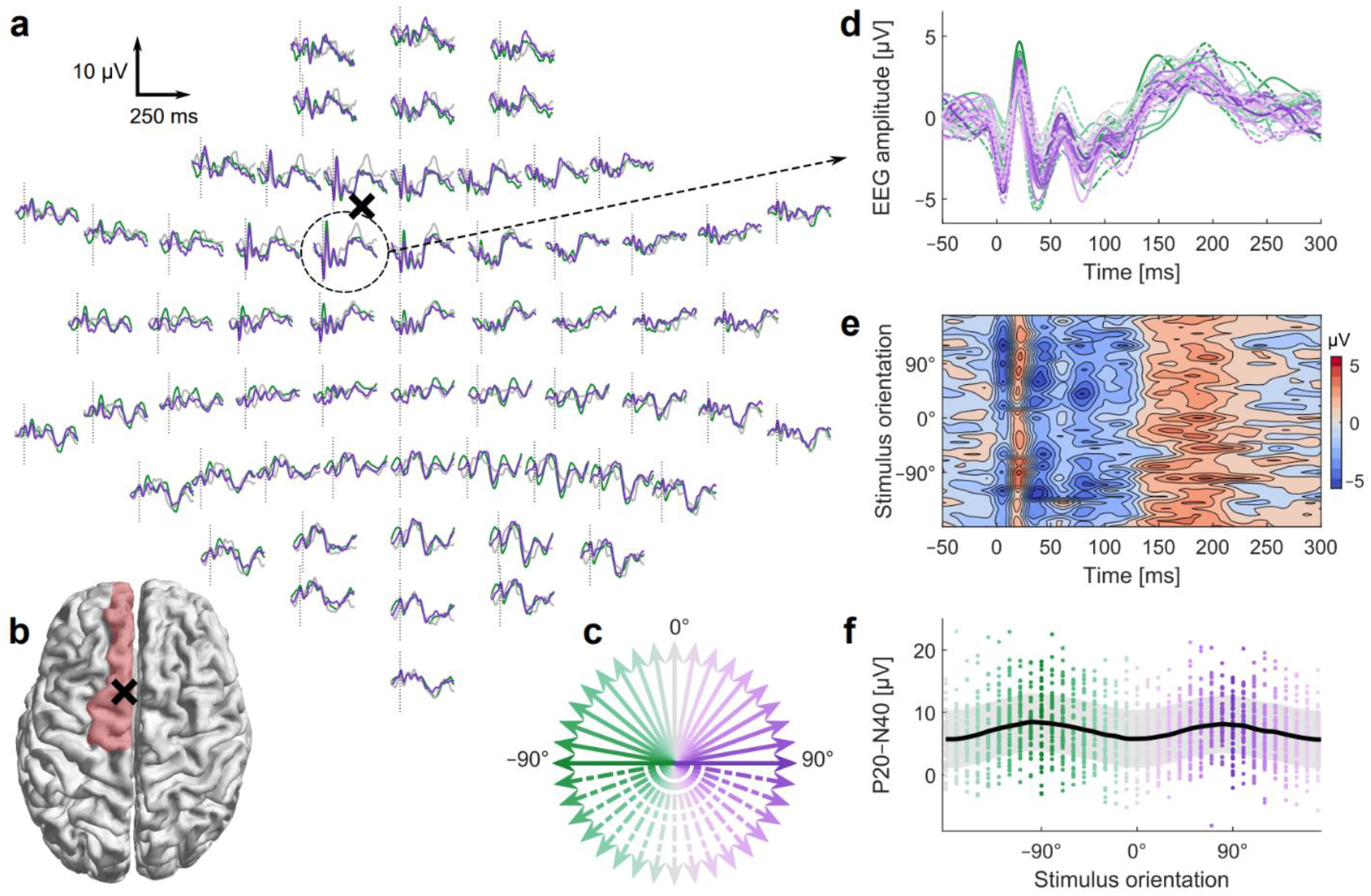
Orientation dependency of the TMS-evoked EEG responses on the left pre-SMA (Subject 1). **a** Time courses of the TEPs in all channels with selected stimulation orientations (−90°, 0°, 90°). The black cross marks the position of the transducer center relative to the electrode locations (**a**) and the stimulation site relative to the brain anatomy (**b**). In **b**, the left superior frontal gyrus is highlighted in red. **c** The 36 stimulation orientations. The colours and line styles of the arrows indicate the corresponding stimulation orientations in **a** and **d**. **d** Enlarged TEP time courses of the channel FC1 with all stimulation orientations. **e** Isocolour plot of the TEP time courses with different stimulus orientations in FC1. **f** Peak-to-peak amplitudes of the P20−N40 complex in FC1 with different stimulation orientations. The dots depict the single-trial responses, the solid trace is a mean curve, and the shaded area illustrates the standard deviation.

Figure 2d visualizes the time course of the TEPs in channel FC1 sited near the stimulation location. The first deflections after the stimulus onset peaked at around 10 ms (negative), 20 ms (positive deflection, named P20), and 40 ms (negative, N40). The amplitudes of these early components depended on the stimulation orientation as clearly seen in Fig. 2e. In addition, the peak-to-peak amplitude of the P20−N40 complex varied as a function of the stimulation orientation in a sinusoidal way, having maxima at around −90° and 90° (see Fig. 2f), when the reference orientation 0° was in the posterior-to-anterior direction. The orientation dependency of the P20−N40 amplitude with all six subjects is presented in Fig. 4a–f.

### Experiment 2: automated EEG-based orientation search

Figure 3 presents two examples of the automated EEG-based orientation search with Subject 1. The first example (Fig. 3a–c) displays how the search rapidly converged to one of the maxima of the P20–N40 response curve while the second example (Fig. 3d–f) demonstrates an additional switch from one maximum to another one during the search process. Figure 3c,f reveals that the sampling was guided in such a way that about the first ten samples were almost evenly distributed across the orientations, whereas the rest of the samples were mainly centred around the current estimate of the optimal orientation. The example search outcomes are visualized in Fig. 3a,d, which also illustrates the large variation of the single-trial responses.

**Figure 3.**
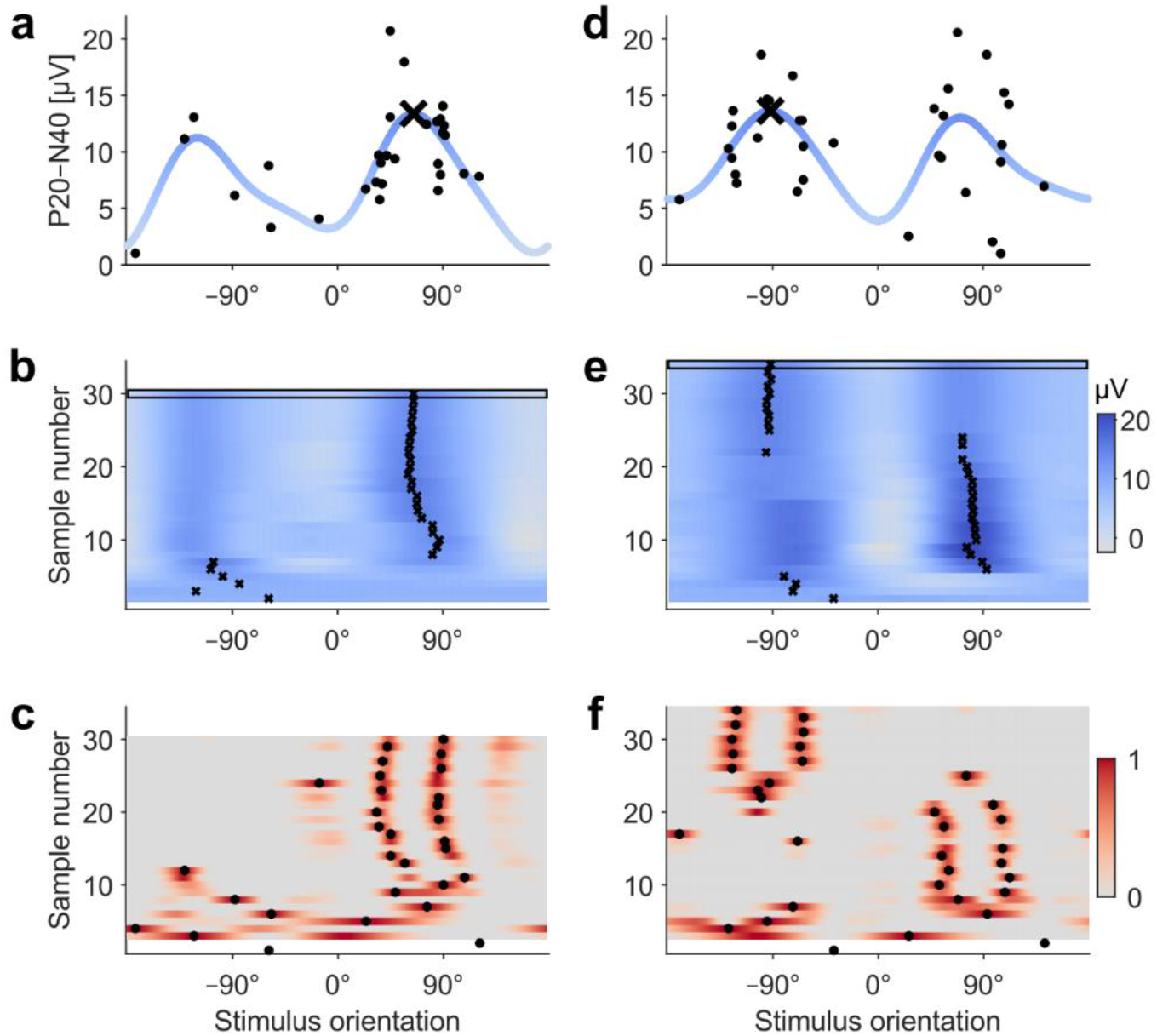
Two examples (**a–c** and **d–f**) of the EEG-based orientation search (Subject 1). **a,d** The search outcome, i.e., the found optimal orientation is marked with a black cross. The acquired single-trial P20–N40 amplitudes are presented with black dots. The blue trace illustrates the final posterior mean curve (modelled behaviour of the response curve). **b,e** The progress of the estimated optimal orientation during the search run. The posterior mean curves computed based on the gathered P20–N40 responses are encoded with coloured rows (grey to blue), and the black crosses indicate the estimated optimal orientation (maximum of the posterior mean curve) on each iteration. The uppermost row (indicated with a black rectangle) correspond to the blue posterior mean curve in **a** and **d**. **c,f** The sampling order. After two randomly sampled orientations (with a 180° difference), we sampled the orientation where the knowledge-gradient function (grey-to-red-coloured rows) reached its maximum (black dots).

The ability of the closed-loop search to find the optimal stimulus orientation is visualized in Fig. 4, which illustrates how the single search outcomes (optimized stimulus orientation) grouped around the ground-truth orientations (maxima of the mean curves determined in Experiment 1). The convergence of the single search runs is presented in Fig. 4g. The average accuracy of the EEG-based orientation search over the 120 repetitions was 18° (individual average accuracies 5–43°; median 10°). When excluding Subject 3, whose EEG data were excessively noisy, the average accuracy was 13° (median 9°). This 13° deviation from the ground truths corresponds on average to a 0.25-μV or 3% decrease in the amplitudes of the P20– N40 mean curve maxima. Eighty-eight per cent of the optimization results were closer than 25° to the ground truth (77% when including Subject 3). The average number of TMS pulses needed in the search was 42 (range 33–48 among the subjects; median 37). The accuracy of the estimated optimal orientation and the number of pulses needed in the search tended to depend on the signal-to-noise ratio (SNR) of the optimized TEP feature as can be seen in Fig. 4h–i: the better the SNR, the less error there was in the search outcomes and the smaller number of stimuli needed for convergence.

**Figure 4.**
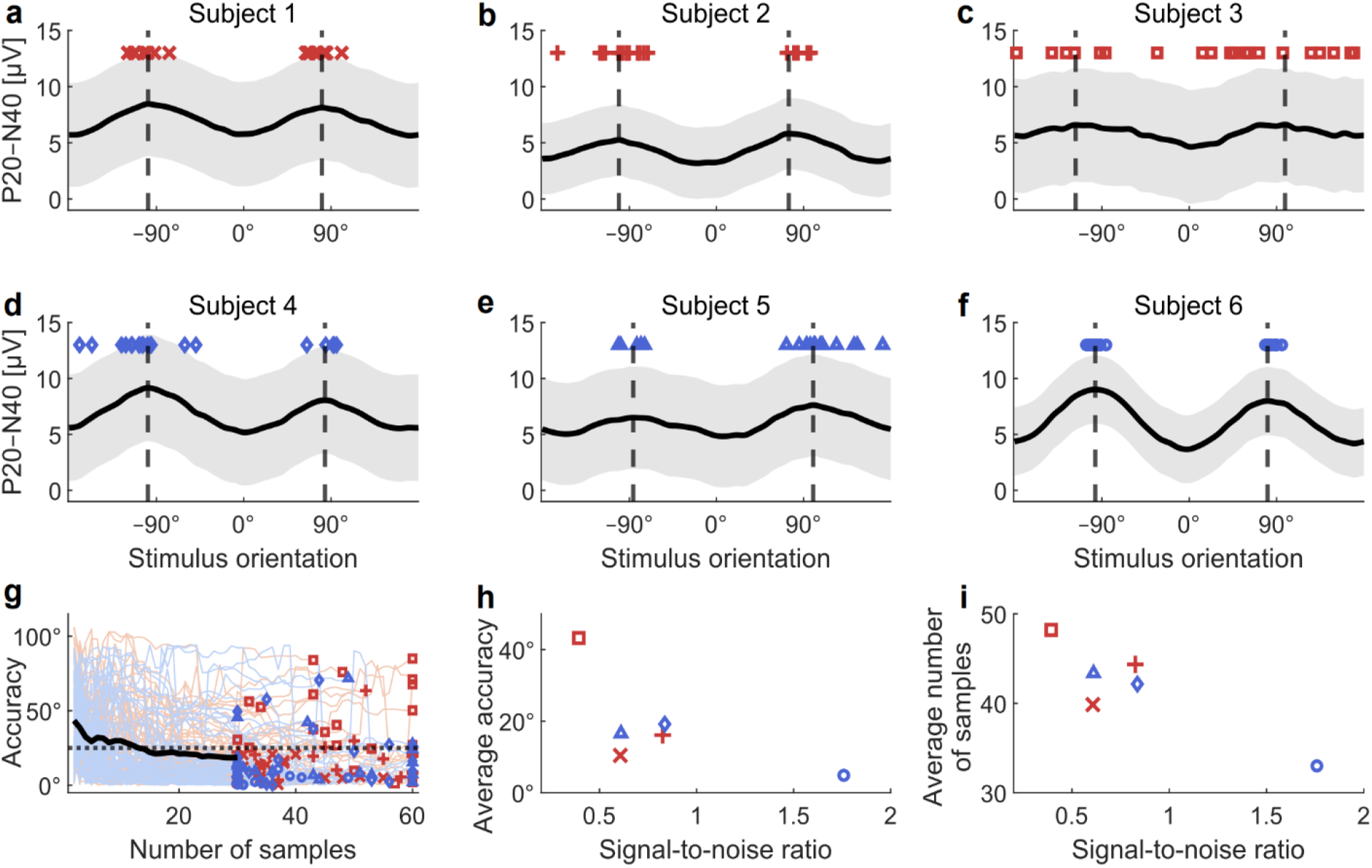
Results of the validation of the EEG-based orientation search. **a–f** Subject-wise search results. Optimized orientations from single searches are depicted with red and blue markers. Mean curves of the optimized P20–N40 amplitudes constructed from the data measured in Experiment 1 are visualized with solid black lines, and the vertical dashed lines illustrate the maxima of the mean curves (ground-truth optimal orientations). Shaded grey areas indicate the standard deviation of the single-trial P20–N40 amplitudes. Subjects of Group A are presented in **a–c** and subjects of Group B in **d–f**. **g** Convergence of the automated orientation searches. Red (Group A) and blue (Group B) lines depict the convergences of the single search runs, and the black curve represents the average error until the minimal number of samples (30) is reached. The end results are presented with red and blue markers. The horizontal dotted line marks an error of 25°. **h** Average error in the search results as a function of the signal-to-noise ratio. **i** Average number of samples needed as a function of the signal-to-noise ratio. The marker shapes and colours in **g–i** corresponds the subject-wise marker styles in **a–f**.

## Discussion

The ability to adjust TMS parameters automatically based on online EEG feedback opens new prospects for scientific and clinical applications of TMS. The results of our algorithm performance evaluations (Experiment 2) demonstrate that our automated closed-loop search provides an easy, fast, and user-independent way to determine TMS targets based on evoked EEG signals. This is, to our knowledge, the first time that post-stimulus EEG responses helped to optimize TMS efficacy in a closed-loop manner.

The automated orientation search (Experiment 2) performed well in all subjects except one whose EEG was contaminated by excessive scalp muscle activity present throughout the measurements. The differences in the algorithm performance are explained by the alterations in the SNR of the data: better SNR increases the accuracy (Fig. 4h) and reduces the number of required iterations in the search process (Fig. 4i). The accuracy of the optimized orientations would likely get better by increasing the number of iterations in the search, at the cost of increasing the search time. Depending on the application, different criteria can be chosen to terminate the search.

Even though the EEG-based orientation search worked well in most cases, there are a few erroneous search results visible in Fig. 4a–g. These outliers resulted from the fact that single-trial TEPs, and thus the extracted P20−N40 amplitudes, are very variable (see example distributions in Figs. 2f and 3a,b). This variation makes the optimization challenging, as sometimes stimulation with the optimal orientation may evoke small P20−N40 amplitudes, and, with non-optimal orientation, the amplitudes can be large by chance. The changes in the single-trial EEG responses with different stimulation orientations are difficult or even impossible to register and interpret visually by the operator. However, our user-independent algorithm uncovers the optimal orientation often even with a few tens of pulses in different directions. This is remarkably few in contrast to traditional TMS–EEG in which several tens or even a few hundred trials with the same stimulation parameters are averaged together for the offline analysis and interpretations. The performance of our closed-loop algorithm shows that even a single-trial TEP contains useful information when combined across different TMS parameters in a meaningful way.

The EEG feature we selected for the optimization was the amplitude of the P20–N40 complex, as it showed apparent orientation dependency in Experiment 1 across the Group A subjects (algorithm test data). The amplitudes of the early components (within the first 50 ms after the pulse) have been used also in other studies to adjust TMS parameters [28,29], as the early TEP deflections are thought to reflect cortical excitability [29,48–50]. Furthermore, peak-to-peak amplitudes are less susceptible to baseline drifting than the amplitudes of single peaks. In this study, we applied the automated search for the peak-to-peak amplitudes in one channel, namely FC1. However, there are many other aspects in the EEG signal that could be optimizable features in closed-loop TMS–EEG. For example, one could employ the signal from several channels in form of a spatial filter [32] to capture the signal from a desired cortical area or utilize the multi-channel EEG to follow and optimize the signal propagation between different cortical areas.The EEG feature we selected for the optimization was the amplitude of the P20– N40 complex, as the TEP deflections within the first tens of milliseconds after the pulse are thought to reflect cortical excitability at least in the primary motor cortex [48]. Furthermore, peak-to-peak amplitudes are less susceptible to baseline drifting than the amplitudes of single peaks. In this study, we applied the automated search for the peak-to-peak amplitudes in one channel, namely FC1. However, there are many other aspects in the EEG signal that could be optimizable features in closed-loop TMS–EEG. For example, one could employ the signal from several channels in form of a spatial filter [32] to capture the signal from a desired cortical area or utilize the multi-channel EEG to follow and optimize the signal propagation between different cortical areas.

We tested the automated orientation search on pre-SMA since it is often easy to acquire high-quality artifact-free TMS–EEG data from that area [51]. In addition, pre-SMA is relevant when studying, for example, cognitive control [52] or motor learning [53] with TMS; it has been studied as a potential target for treating patients with obsessive-compulsive disorder and essential tremor [2]. We presume that this kind of orientation search would work also on other brain areas, but it would require further systematic mapping to find out what are the suitable time intervals for detecting the peak amplitudes, as the number of TEP deflections and their latencies vary across the brain [18]. Other stimulation sites, especially lateral ones, may be more prone to muscle artifacts and require sophisticated artifact removal methods to ensure reliable functioning of closed-loop TMS–EEG algorithms. As in this work we stimulated in all directions, we unsurprisingly observed some muscle artifacts with certain stimulation directions with two subjects (1 and 6) when visually inspecting the averaged raw signals from Experiment 1. The muscle artifacts were relatively small (maximum amplitude 20–30 μV) and appeared only in a few channels (up to four) lateral or anterior to the channel of interest (FC1). The SOUND algorithm applied offline for the data of Experiment 1 diminished these muscle artifacts; the artifacts were, however, not necessarily completely removed. Despite possible residual artifacts with some of the subjects, the observed sinusoidal orientation dependency in the amplitude of the P20–N40 complex was consistent across all subjects (Fig. 4a–f). Therefore, we believe that the observed effect originates from the brain and not from artifacts. The online preprocessing in Experiment 2 included no artifact cleaning. Therefore, some of the single-trial responses of Subjects 1 and 6 contained minor TMS-related muscle artifacts in channels other than the channel of interest (FC1). All search outcomes of these subjects were close to the ground truth optima determined from SOUND-cleaned data (see Fig. 4a,f); thus, muscle artifacts seem not to have affected the performance of the closed-loop optimization.

In addition to the stimulation orientation, our algorithm could optimize EEG responses as a function of other stimulation parameters, such as the stimulation location, intensity, or timing (with respect to ongoing brain activity), or all of them simultaneously. In this work, we determined the stimulation location and intensity manually based on visual inspection of averaged TEPs [29]. We believe that the whole procedure can, however, be automated by combining the presented closed-loop algorithm, for example, with a 5-coil mTMS system that allows adjusting the stimulation location and orientation without coil movement [54]. An advanced algorithmic implementation may thus make visual evaluation and manual assessment redundant in the future. Such an implementation needs, however, to be highly reliable to avoid erroneous results. Basically, the presented automated approach is suitable for optimizing any feature derived from EEG with respect to one or more parameters.

We controlled the stimulation orientation with a two-coil transducer comprising two simultaneously operated overlapping figure-of-eight coils. This provides an easy and effortless adjustment of the stimulation orientation, as it requires no manual movement of the transducer. The EEG-based target automation presented here could be implemented with a robot-controlled TMS system or with manual coil placement. However, the physical coil shift takes time, and the movement of the coil between the consecutive pulses causes motion artifacts in EEG data that require a few additional seconds to get stabilized, increasing the search time.

The systematic mappings of TMS–EEG responses as a function of the stimulation orientation in Experiment 1 revealed that the amplitudes of the TEP peaks depend on the stimulation direction on the pre-SMA in a sinusoidal way. There are a few examples in the literature showing that the stimulation orientations 45° or 90° apart result in dissimilar TEPs on the primary motor cortex [15] and occipital areas [16]. Thus, it was foreseeable to observe an orientation dependency, but this is the first study showing examples of the behaviour of the TEP responses when varying the stimulation orientation systematically in all directions. The optimal stimulus orientations that produce maxima of the P20–N40 curves were reached with E-fields directed approximately along the medial–lateral direction. These directions are perpendicular to the global orientation (posterior–anterior) of the targeted superior frontal gyrus (Fig. 2b). This is in line with the behaviour of motor responses when stimulating the motor cortex [6] and the E-field modelling on other areas [55], which have shown that TMS is most effective when the stimulating E-field is directed normal to the sulcal walls. Additionally, we found that the sinusoidal form of the TEP amplitude curve supports the cosine model for TMS [56], which suggests that the TMS effect in any part of the cortex depends on the cosine of the angle between the cortical column orientation (normal to the cortical surface) and the direction of the E-field. E-field can be precisely oriented based on anatomy with the help of existing neuronavigation systems. There are, however, tight curvatures in the folded structures of individual cortices, necessitating stimulus guiding with neurophysiological feedback such as EEG.

We anticipate that the presented closed-loop optimization maximizing the TMS effect based on EEG signal would increase the comparability of TMS research by decreasing the variation in experimental designs. Targeting based on the amplitudes of the early TEP deflections resembles the motor-response-based selection of stimulus location and orientation (often called hotspot search) in the primary motor cortex. In contrast, EEG-based guiding enables setting the stimulation parameters with neurophysiological feedback in any part of the superficial cortex. Being able to analyse EEG signals online and utilize them to adjust TMS delivery enables guiding TMS therapies so that the desired plastic effects are maximized. This is an indispensable step towards making TMS a more effective clinical tool.

## Conclusions

We demonstrated that it is possible to adjust the stimulation orientation based on EEG signals effortlessly with multi-locus TMS and showed that even single-trial TEPs contain features that can be optimized. The presented method provides a way for automated and individualized TMS targeting with neurophysiological feedback over a large part of the cortex. Moreover, closed-loop TMS–EEG holds great promise for making TMS treatments more effective.

## Acknowledgements

We thank Baran Aydogan for helping with the acquisition of the MRIs, Science-IT at Aalto University School of Science for the computational resources, Mikko Raskinen (Aalto University) for the photos, and Prof. Massimini’s group at the University of Milan for providing Matlab tools for generating noise-masking audio tracks and visualizing TMS–EEG responses.

